# JanusX: an integrated and high-performance platform for scalable genome-wide association studies and genomic selection

**DOI:** 10.64898/2026.01.20.700366

**Authors:** Jingxian Fu, Anqiang Jia, Haiyang Wang, Hai-Jun Liu

## Abstract

As genomic datasets expand in both sample size and marker density, genome-wide association studies (GWAS) and genomic selection (GS) require workflows that remain statistically rigorous, computationally efficient, and reproducible across the full analysis path, from genotype matrix to decision-relevant outputs. Here we present JanusX, an integrated high-performance framework that provides a streamlined, user-oriented workflow for GWAS and GS by unifying data handling, model execution, and visualization. Across simulated and real datasets, JanusX maintained high concordance with established baselines while substantially reducing runtime and memory usage. In GWAS, JanusX achieved up to a 19-fold speedup over GEMMA in linear mixed model (LMM) inference, and implemented additional LMM inference based on a sparse genomic relationship matrix with GRAMMAR-Gamma calibration, alleviating computational and memory bottlenecks in large-scale cohorts. JanusX also provides a FarmCPU implementation within its GWAS module, achieving a median 11.4-fold runtime improvement and reducing peak memory usage by 84.9% relative to rMVP. In GS, JanusX integrates an optimized best linear unbiased prediction (BLUP) backend that adaptively selects sample- and SNP-space solvers and incorporates a Preconditioned Conjugate Gradient (PCG) solver. This implementation efficiently completes five-fold cross-validation of 500k individuals × 500k single-nucleotide polymorphisms (SNPs) in 35.1 minutes with only 14.3 gibibyte (GiB) of peak memory. Beyond BLUP, JanusX integrates Bayesian and machine-learning predictors under a single interface with compact automatic tuning to ensure robust cross-model performance. JanusX therefore enables efficient locus discovery and genomic prediction under consistent analytical assumptions, even in large-scale cohorts.

## Introduction

Genome-wide association studies (GWAS) are central to dissecting the genetic architecture of complex traits (Huang and Han, 2014, Visscher *et al*., 2017), whereas genomic selection (GS) is increasingly used to predict genetic merit and support breeding decisions (Meuwissen *et al*., 2001, Georges *et al*., 2019). These two analytical paradigms are now frequently applied to the same breeding populations but typically rely on distinct and fragmented software ecosystems. As cohort sizes and marker densities continue to increase (Kurki *et al*., 2023, Loya *et al*., 2025), this fragmentation complicates consistency across key processes such as preprocessing, model execution, result comparison, and visualization. These challenges, in turn, increase computational requirements, memory demand, and reproducibility risks (Davis-Turak *et al*., 2017).

Computational constraints are particularly acute for mixed-model analyses. In GWAS, linear mixed models (LMMs) remain a standard approach for controlling population structure and cryptic relatedness through genomic relationship matrices (GRMs) (Yu *et al*., 2006), and BLUP-family models are foundational for GS (VanRaden, 2008, Endelman, 2011, Covarrubias-Pazaran, 2016). Algorithmic advances such as dense-GRM eigenvalue decomposition (EVD) (Lippert *et al*., 2011, Zhou and Stephens, 2012), sparse-GRM factorization (Jiang *et al*., 2019), fixed-variance scanning (Kang *et al*., 2010), low-rank representations (Lippert *et al*., 2011), GRAMMAR-Gamma calibration (Svishcheva *et al*., 2012) and iterative solvers (Loh *et al*., 2015) have greatly improved scalability. Nevertheless, these advancements are often confined to specialized tools that differ significantly in input formats, model interfaces, output conventions and downstream visualization capabilities. As a result, researchers often face a trade-off between high-performance tools optimized for specific tasks and fragmented toolchains required for complete analyses, which exposes them to steep learning curves, reproducibility risks, and growing maintenance challenges as datasets increase in scale.

Here, we present **JanusX**, an integrated high-performance platform for GWAS and GS, with supporting modules for population-structure analysis and post-analysis visualization. JanusX is designed to accommodate genetic and breeding studies, ranging from medium-sized experimental populations to large cohorts, by addressing three central objectives: 1) providing a unified workflow spanning data handling, model execution, and visualization; 2) maintaining statistical rigor and ensuring high concordance with established baselines across GWAS and GS models; and 3) improving scalability through optional dense- and sparse-GRM backends, iterative solvers, and system- and implementation-level optimizations. Together, these features make integrated locus discovery and genomic prediction more computationally feasible, reproducible, and accessible across diverse studies.

## Results

### Overview of JanusX

JanusX provides a unified workflow for GWAS and GS, starting from harmonized genotype-phenotype inputs and producing standardized outputs, including statistical summaries, model-evaluation results, and visualization outputs (Fig. 1A). To streamline analysis, JanusX automatically decodes genotype, phenotype, and optional covariate inputs, applies quality control, and matches samples across these data types to generate an analysis-ready dataset. This design eliminates redundant format conversions and minimizes cross-tool sample-matching errors. When required by downstream analyses, shared analytical representations are generated directly from genotype data. For example, GRMs are constructed using generalized matrix-matrix multiplication (GEMM), and principal components (PCs) are computed through either EVD or randomized singular value decomposition (rSVD). The harmonized dataset is then routed into either association-mapping or genomic-prediction workflows. The association-mapping module supports a wide range of models, including the general linear model (‘-lm‘), LMM (‘-lmm‘), fixed-variance LMM (‘-fvlmm‘), sparse-GRM LMM (‘-splmm‘), and FarmCPU (‘-farmcpu‘). Association statistics were evaluated using Wald and likelihood-ratio (LR) testing. The genomic-prediction module supports BLUP (‘-BLUP‘), BayesA (‘-BayesA‘), BayesB (‘-BayesB‘), BayesC (‘-BayesCpi‘), and various machine learning models for genomic estimated breeding value (GEBV) prediction, along with build-in model evaluation.

**Figure 1.**
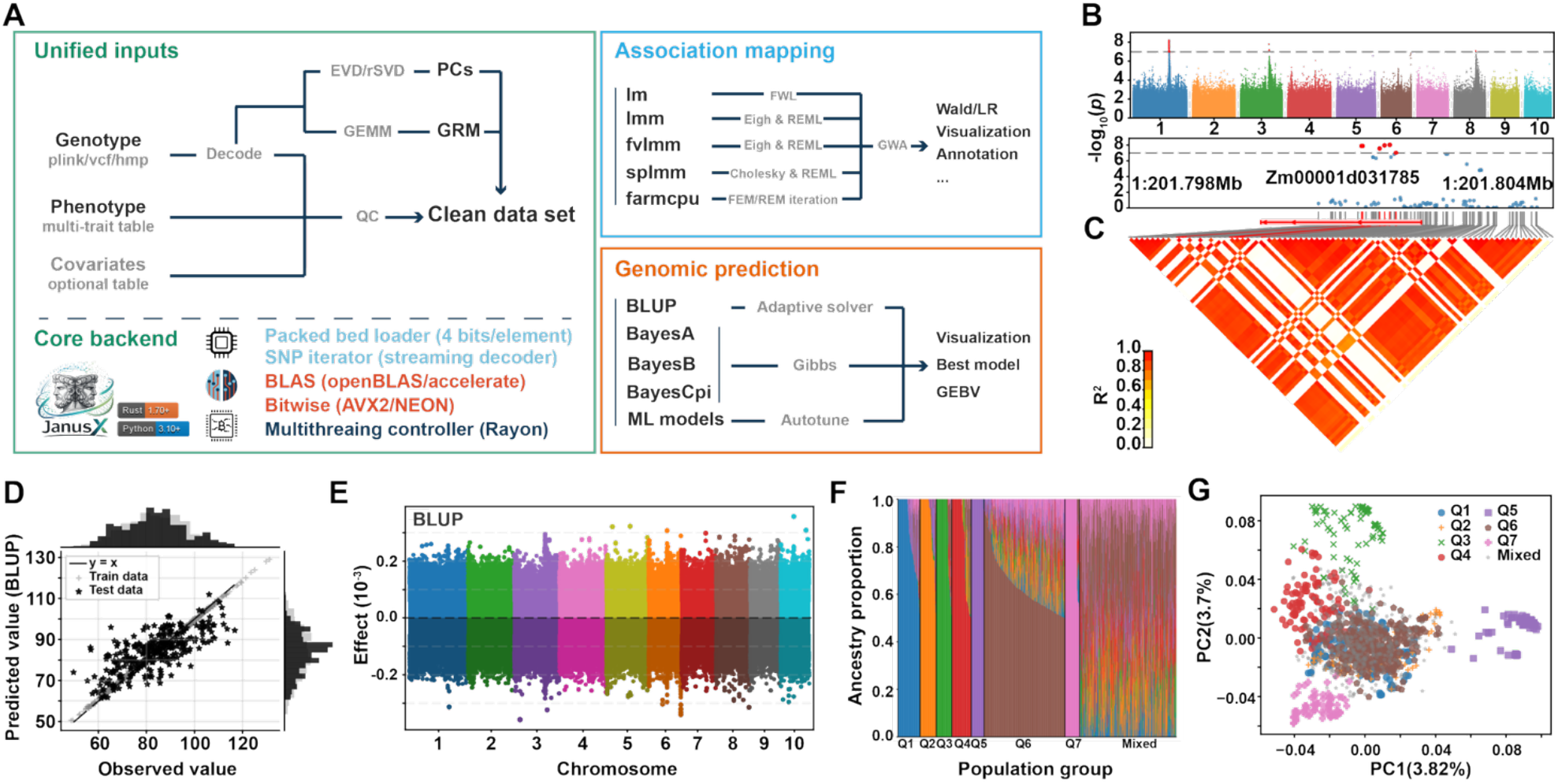
The JanusX framework architecture and visualization outputs. **(A)** Schematic of the unified computational workflow for GWAS and GS. Unified genotype, phenotype, and optional covariate inputs are converted into shared analysis-ready representations and directed into association-mapping and genomic-prediction modules. The backend provides hardware-aware acceleration through features like packed-BED loading, SNP-iterator streaming, BLAS-accelerated matrix computation, bitwise operations, and multithreaded execution. In association mapping panel, notations such as FWL, REML, Cholesky, FEM and REM denote Frisch-Waugh-Lovell residualization, restricted maximum likelihood, Cholesky decomposition, fixed-effect model, and random-effect model, respectively. **(B, C)** Representative outputs from the GWAS module, including a Manhattan plot for genome-wide association signals and a triangular LD block plot for local linkage disequilibrium (LD) structure. **(D, E)** Representative outputs from the GS module, including a scatter plot of predicted versus observed phenotypes and a Manhattan-style plot of marker-effect estimates. **(F, G)** Representative outputs from the ‘fastpop‘ and PCA modules for population-structure analysis, including an ancestry-proportion stacked bar plot and a PCA scatter plot colored by inferred population groups.

JanusX also provides built-in visualization modules for both GWAS and GS outputs (Fig. 1B-E and Supplemental Note 1). For GWAS, JanusX generates Manhattan plots for marker-level association statistics (Fig. 1B) and linkage disequilibrium (LD) block triangular plots for local haplotype or LD structure (Fig. 1C). For GS, JanusX provides model evaluation plots for predictive performance assessment (Fig. 1D) and Manhattan-style plots for estimated marker effects or feature importance scores (Fig. 1E). In addition, JanusX integrates ‘fastpop‘, a population-structure analysis module optimized within the general analytical framework of ADMIXTURE (Saurina-i-Ricos *et al*., 2026). The resulting ancestry proportions can be directly visualized as stacked bar plots (Fig. 1F), and population assignments can be combined with genotype-derived PCs to generate principal components analysis (PCA) scatter plots (Fig. 1G).

We therefore focused the following benchmarks on JanusX modules that implement internally optimized backends, primarily the LMM-family GWAS methods and the adaptive BLUP backend for GS. We first assessed their concordance with established baseline tools and then quantified runtime and memory performance for the corresponding optimized workflows. FarmCPU for GWAS and Bayesian models for GS were included as additional real-data benchmarks, whereas machine-learning predictors were excluded from the core performance benchmarks because they rely on wrappers around mature external Python libraries (Pedregosa *et al*., 2011) rather than being native JanusX implementations.

### Benchmarking the GWAS module

The GWAS benchmarks focused on JanusX’s association backends designed to support various analytical scales and model configurations. JanusX-lmm implements a dense-GRM full LMM strategy based on EVD (Fig. 2A), and re-estimates variance components for each SNP, providing a high-fidelity comparison point similar to GEMMA-lmm (Zhou and Stephens, 2012). JanusX-fvlmm uses the same LMM framework but estimates variance components once under the null model and keeps them fixed during genome-wide scanning. For large-cohort analyses, JanusX-splmm implements sparse-GRM LMM inference with optional GRAMMAR-Gamma calibration (Fig. 2A), analogous to GCTA-fastGWA (Jiang *et al*., 2019). In parallel, JanusX provides a FarmCPU implementation (Liu *et al*., 2016), which was benchmarked against rMVP-FarmCPU (Yin *et al*., 2021) using real maize CUBIC datasets.

**Figure 2.**
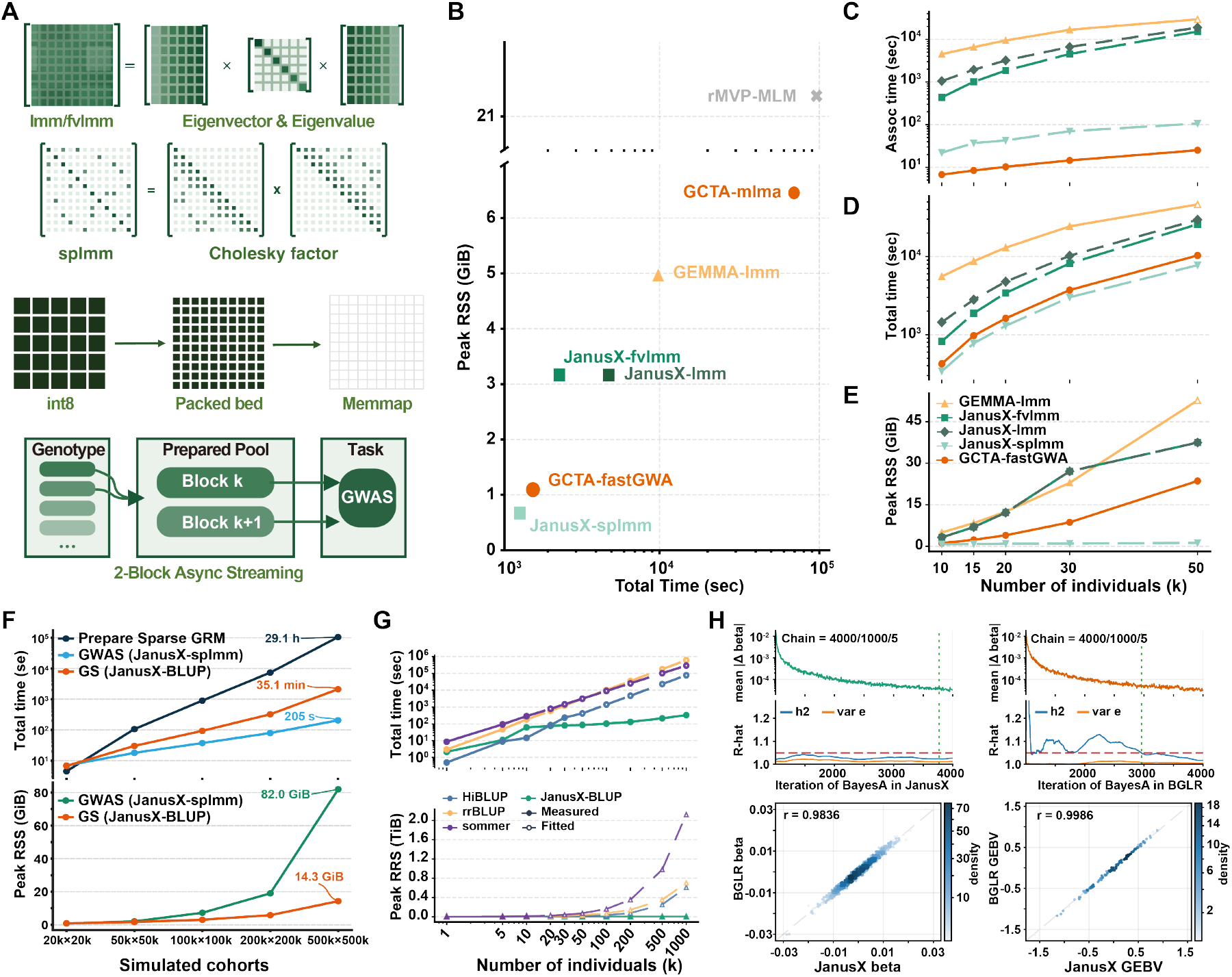
Backend optimizations and benchmarks of GWAS and GS module in JanusX. **(A)** Overview of JanusX backend optimization. Dense-GRM EVD supports ‘-lmm‘ and ‘-fvlmm‘ in small-to medium-sized cohorts, whereas Cholesky-based sparse-GRM factorization optimally supports ‘-splmm‘ in large cohorts. Memory-efficient methods, including packed-BED representation, memory-mapped access, streaming decoding, and two-block asynchronous buffering, significantly reduce memory pressure and accelerate genotype throughput. **(B)** Single-core runtime and peak resident set size (RSS) comparison of mixed-model GWAS tools on a simulated benchmark dataset. **(C-E)** Sample-size scalability of GWAS backends, measured by association-stage runtime (C), end-to-end runtime including GRM construction (D), and peak RSS (E). Association-testing time (Assoc Time) and “Total Time” denote the runtimes for the association testing phase and the entire GWAS workflow, respectively. **(F)** Biobank-scale stress test of JanusX GWAS and GS modules on simulated genotype matrices accounting for matched numbers of individuals and SNPs. **(G)** Sample-size scalability of BLUP-family GS workflows, showing total runtime and peak RSS across increasing sample sizes. **(H)** Concordance of BayesA between JanusX and BGLR, with diagnostics summarizing convergence of posterior marker effects and variance components. Chain labels indicate the combination of total MCMC iterations, burn-in iterations, and thinning intervals. Density-colored scatter plots show the agreement in posterior marker-effect estimates and GEBVs, with points color-coded by local density. R-hat is the Gelman-Rubin diagnostic, which evaluates chain convergence (values close to 1 signify reliable convergence). Dashed lines mark the burn-in cutoffs and the R-hat threshold of 1.05.

We first assessed whether these implementations matched established reference tools before runtime and memory performance evaluation. Concordance was evaluated using both real genotype-phenotype datasets and simulated phenotypes generated from empirical genotype panels. The real-data benchmark used maize CUBIC dataset (Liu *et al*., 2020), whereas simulation datasets used empirical genotype matrices from the maize CUBIC and RiceAtlas panels (Ma *et al*., 2025). Simulations preserved authentic minor allele frequency (MAF) spectra, LD patterns, and population structures (Fig. S1), facilitating benchmarks against tools like GEMMA-lmm and GCTA-fastGWA (LMM-family concordance), and rMVP-FarmCPU (FarmCPU concordance).

For both the 12 simulated genetic-architecture scenarios described in Methods and five representative real traits from the maize CUBIC population, concordance was evaluated among the top 1% most significant markers to avoid agreement being dominated by null background SNPs. JanusX-lmm and JanusX-fvlmm showed near-perfect concordance with GEMMA-lmm (Fig. S2A), achieving Pearson correlation coefficients (r) above 0.9991 and Spearman correlations (ρ) for association significance above 0.9844 (Tables S1 and S3). Sparse-GRM LMM results from JanusX-splmm aligned strongly with GCTA-fastGWA (Fig. S2B), with Pearson correlations for marker effects ranging from 0.9993 to 1.0000 and Spearman correlations for association significance ranging from 0.9804 to 1.0000 (Tables S2 and S4). Genomic inflation factor (λ_GC_) estimates from JanusX-splmm were slightly lower than those from GCTA-fastGWA, indicating slightly more conservative calibration (Tables S2). FarmCPU results demonstrated complete pseudo-QTN agreement with rMVP-FarmCPU across 23 maize CUBIC traits (Fig. S2C). These results confirm that the optimized GWAS implementations in JanusX preserve high concordance with established baseline tools across simulated and real-data settings.

To place the JanusX LMM-family GWAS backends in the context of widely used mixed-model association tools, we first performed a single-core benchmark on a simulated dataset containing 10,000 individuals and 1,000,000 SNPs generated with the JanusX simulation module. The comparison included JanusX-lmm, JanusX-fvlmm and JanusX-splmm, together with GEMMA-lmm (Zhou and Stephens, 2012), GCTA-mlma (Yang *et al*., 2011), GCTA-fastGWA (Jiang *et al*., 2019), and rMVP-MLM (Yin *et al*., 2021). All tools were run using a single CPU core. JanusX-splmm achieved the best overall efficiency, completing the full workflow in 22.3 minutes with a peak memory usage (resident set size, RSS) of only 0.67 GiB (Gibibyte, binary; 1 GiB = 1024 MiB = 2^30^ bytes). By comparison, GCTA-fastGWA required 26.9 minutes with 1.09 GiB, while GEMMA-lmm required 2.72 hours and 4.97 GiB (Fig. 2B). Further parallel scaling tests (2-32 cores) demonstrated strong performance for JanusX-lmm and JanusX-splmm, achieving up to a 30.2-fold runtime improvement for sparse-GRM workflows, outperforming conventional tools GEMMA-lmm and GCTA-fastGWA (Fig. S3A).

The advantage of JanusX-splmm persisted in scalability analyses against GCTA-fastGWA (Fig. 2C-E and Fig. S3B). Dense-GRM LMM methods, including GEMMA-lmm, JanusX-lmm, and JanusX-fvlmm, were included as reference baselines, but their computational expense sharply escalated with increasing dataset size, particularly as the number of samples increased. For datasets spanning 1-20 million SNPs at 10,000 individuals, and 10,000-50,000 individuals at 1 million SNPs, JanusX-splmm consistently achieved lower end-to-end runtime and peak memory usage in the total GWAS workflow compared to other methods. In the SNP-scaling benchmark, for example, JanusX-splmm reduced total runtime from 8533.3 seconds (GCTA-fastGWA) to 6,676.7 seconds and peak memory from 3.55 GiB to 1.53 GiB at 20 million SNPs (Fig. S3B). Similarly, in the sample-size scaling benchmark, JanusX-splmm reduced runtime from 10,322.1 seconds to 7,751.6 seconds and peak memory usage from 23.59 GiB to 1.21 GiB at 50,000 individuals (Fig. 2D, E). Although GCTA-fastGWA demonstrated faster association-testing phases (Fig. 2C), its computational overhead during GRM construction negated this advantage, leading to longer overall runtime and higher memory consumption in the total GWAS workflow.

These efficiency gains extended beyond LMM-based association testing. We benchmarked FarmCPU against 23 agronomic traits using the maize CUBIC genotype panel. Compared to rMVP-FarmCPU, JanusX-FarmCPU achieved dramatic resource reductions, requiring only 9.5% ± 2.6% of the runtime and 14.5% ± 0.8% of the peak RSS. Notably, JanusX-FarmCPU maintained complete agreement with rMVP-FarmCPU in the pseudo-QTNs selected through FEM-REM iterations (Fig. S2C).

To further assess whether the JanusX GWAS module can handle biobank-scale genotypic dimensions, we conducted an ultra-large capacity stress test using JanusX-splmm on simulated random genotypes. At the largest tested scale of 500,000 individuals and 500,000 SNPs, JanusX-splmm completed the association scan in 205 seconds after sparse-GRM construction (Fig. 2F). Sparse-GRM construction dominated the total runtime, while the marker-scanning step demonstrated consistent efficiency. These results show that JanusX can execute GWAS seamlessly at biobank-scale dimensions, shifting the primary computational bottleneck to sparse-GRM preparation.

### Benchmarking the GS module across Bayesian and BLUP models

To evaluate the GS module in JanusX, we first benchmarked JanusX-BLUP against established tools, including rrBLUP (Endelman, 2011), sommer (Covarrubias-Pazaran, 2016), HIBLUP (Yin *et al*., 2023) and JanusX Bayesian prediction models against BGLR (Perez-Rodriguez and de los Campos, 2014). JanusX-BLUP produced near-identical GEBV predictions to rrBLUP, HIBLUP, and sommer, with pairwise Pearson correlations ranging from 0.9994 to 1.0000 (Fig. S4A). For Bayesian models, we performed thorough convergence diagnostics to determine stable sampling lengths for BayesA, BayesB, and BayesC as 4,000, 8,000, and 2,000 iterations, respectively (Fig. 2H and Fig. S4B, C). Posterior marker-effect estimates obtained from JanusX exhibited excellent concordance with BGLR across these models (Fig. 2H and Fig. S4B, C).

To evaluate the computational performance, we conducted stress tests for BLUP-family models using fully simulated genotype and phenotype data generated by the JanusX ‘sim‘ module. At a fixed marker density of 10,000 SNPs, JanusX-BLUP scaled efficiently from 1,000 to 1 million individuals, with total runtime increasing from 2.15 seconds to 330.63 seconds and peak RSS staying below 4.5 GiB (Fig. 2G). In contrast, HIBLUP, rrBLUP, and sommer showed prohibitive memory requirements and failed at larger sample sizes. At a fixed sample size of 5,000 individuals, JanusX-BLUP handled growing marker density seamlessly, with runtime increasing only from 10.92 seconds at 5,000 SNPs to 24.82 seconds at 1 million SNPs and peak RSS remaining under 1.4 GiB (Fig. S5). These results illustrate that the adaptive JanusX-BLUP backend decouples computational load from both sample size and marker density, ensuring scalability even for biobank-scale datasets.

To validate biobank-scale performance, JanusX-BLUP completed a full five-fold cross-validation GS workflow with 500,000 individuals and 500,000 SNPs in 2,105.5 seconds (35.1 minutes) with only 14.3 GiB peak memory (Fig. 2F). This scalability is achieved through adaptive solver selection, where JanusX dynamically transitions to a PCG-based iterative solver under high memory demand, bypassing explicit construction of the full sample-space system.

## Discussion

JanusX demonstrates that workflow fragmentation in large-scale genetic analyses can be effectively overcome through integration with high-performance computation. Its capabilities encompass GWAS, GS, population-structure analysis, and visualization in a unified, scalable framework. By significantly reducing runtime and memory usage for models such as LMMs, FarmCPU in GWAS, and BLUP-family models in GS, JanusX offers an efficient and reproducible pipeline for modern genetic and breeding studies.

Nonetheless, several challenges remain. For GWAS, sparse-GRM construction remained a dominant bottleneck (Fig. 2F), particularly as sample sizes and marker density grow. Future improvements may require distributed or block-wise GRM construction (Lin *et al*., 2024) and marker-reduction strategies (e.g., LD pruning or tag-SNP selection). Sparse-GRM factorization could also be optimized further for structured plant breeding populations. For GS, while Bayesian models show high concordance, large-scale stress testing of Bayesian methods remains limited due to poor parallelizability of conventional MCMC inference (Terenin *et al*., 2020). Developing scalable Bayesian inference techniques should also be a future focus.

## Conclusions and outlook

JanusX resolves the fragmentation inherent in genetic analysis workflows by unifying GWAS, GS, and visualization into a single, reproducible pipeline. Benchmarks verified that JanusX maintains statistical concordance with established tools while dramatically improving computational efficiency. Additional extensions, such as support for multi-locus models (Liu *et al*., 2025), downstream selection strategies, and interactive visualization interface (JanusXweb), will further broaden its applicability and accessibility.

## Methods

### Statistical models for genome-wide association

JanusX implements a diverse suite of statistical models, including general linear model (‘-lm‘), LMM (‘-lmm‘), fixed-variance LMM (‘-fvlmm‘), sparse-GRM LMM with GRAMMAR-Gamma calibration (‘-splmm‘), and FarmCPU (‘-farmcpu‘), all integrated within a unified GWAS backend. These models follow well-established statistical frameworks, prioritizing computational efficiency and streamlined data handling., JanusX optimizes implementation by incorporating low-memory genotype streaming and shared data-management strategies. Detailed formulations and derivations for LMM-family models and FarmCPU are provided in Supplemental Note 2.

### Integrated prediction models for genomic selection

JanusX integrates BLUP-family, Bayesian, and machine-learning predictors under a unified GS interface. For BLUP-family models, JanusX implements GBLUP and rrBLUP, while adapting Bayesian models including hierarchical BayesA, BayesB, and BayesC, following the prior structures implemented in BGLR (Perez-Rodriguez and de los Campos, 2014). Machine-learning models include random forest (RF), extra trees (ET), gradient boosting decision tree (GBDT), extreme gradient boosting (XGB), support vector machine (SVM), and elastic net (ENET), with built-in compact tuning procedures.

For BLUP-family prediction, JanusX provides the ‘-BLUP‘ option and dynamically switches between GBLUP (for sample-space calculations), rrBLUP (for SNP-space calculations), and rrBLUP with a PCG solver based on the relative sizes of the sample and marker datasets. Bayesian predictors employ Gibbs-sampling-based frameworks for marker-effect estimation, while machine-learning predictors are wrapped within a standardized interface for model training, hyperparameter tuning, and evaluation. Further technical details for BLUP-family and Bayesian models are provided in Supplemental Note 3 and Note 4.

### Benchmark real and simulated datasets

JanusX was benchmarked using three distinct dataset classes: real benchmark datasets, empirical genotype panels with simulated phenotypes, and fully synthetic genotype-phenotype datasets. Real datasets were used to assess practical applicability and cross-tool concordance. For GWAS, we used the maize CUBIC panel comprising 1,493 individuals and 8,666,018 SNPs after filtering for MAF ≥ 0.02 and missing rate ≤ 0.05. Four representative agronomic traits were used as case studies for LMM-family GWAS benchmarks, while all 23 measured traits were used for FarmCPU benchmarking. For GS, we used the wheat dataset distributed with BGLR, containing 599 individuals and 1,259 markers after preprocessing. JanusX BLUP-family and Bayesian models were compared with sommer, rrBLUP, HIBLUP, and BGLR under harmonized model settings.

For simulation-based GWAS concordance analyses, phenotypes were generated on empirical genotype backbones from the maize CUBIC and RiceAtlas panels. Causal QTNs were randomly sampled without replacement from filtered variants, with either 10 or 100 QTNs per simulation. Raw causal effects were extracted from Uniform(−1,1) distributions and scaled to match the specified V_QTN_. Polygenic background effects followed *N* (0,1) and were scaled to match the specified V_Bg_. For each simulation condition, three additive variance-component scenarios were modeled where phenotypic variance was partitioned as follows: V_QTN_:V_Bg_:V_E_ = 0.5:0.0:0.5, 0.3:0.3:0.4 or 0.1:0.6:0.3, where V_QTN_, V_Bg_ and V_E_ denote variance explained by causal QTNs, polygenic background and residual environment, respectively.

This produced six simulation scenarios per genotype panel while preserving empirical MAF spectra, LD patterns, genetic marker-density heterogeneity and population structures.

Fully synthetic genotype-phenotype datasets were generated with the JanusX quick-simulation workflow for runtime, throughput, peak memory, and biobank-scale scalability tests. These datasets were used specifically for computational performance evaluations.

### Benchmark computing environments

Performance benchmarks were conducted across varied computing environments according to benchmark scale. Routine GWAS-LMM and FarmCPU benchmarks were executed on Rocky Linux 8.9 (Green Obsidian) cluster nodes equipped with Intel Xeon 8458P processors (3.00 GHz). Tasks were assigned 4 or 8 CPU cores respectively, unless otherwise specified. GS scalability benchmarks, including stress tests with 1,000,000 individuals and 10,000 SNPs, were performed with configurations using up to 32 CPU cores, capped at 64 GiB memory. Biobank-scale stress tests for GWAS and GS, scales up to 500,000 individuals × 500,000 SNPs, were performed on the same platform using 64 CPU cores to evaluate scalability boundaries. For GWAS benchmarks, runtime was assessed as the combined duration of GRM construction and genome-wide marker scanning, excluding visualization steps like plotting. All benchmarked tools were run using their latest stable versions available at the time of analysis (rMVP v1.4.6, GCTA v1.94.1, GEMMA v0.98.5, sommer v4.3.2, rrBLUP v4.6.2, HIBLUP v1.6.0, BGLR v1.1.4, and JanusX v1.0.26) to ensure reproducibility and consistency.

## Data and code availability

All genotype and phenotype datasets used in this study are publicly available as cited. JanusX and benchmark script are available at https://github.com/FJingxian/JanusX.

## Acknowledgements

We would like to thank Liying Feng, Zongyi Sun, Xiaoping Guo, Cang Zhao, Peng Zhang, Yang Wang, Niannian Ma, and Han Lu for helping with extensive testing of the software. This work was supported by the Startup Fund of Huazhong Agricultural University (11020153 to H.-J.L), the Yazhouwan National Laboratory, the Hainan Postdoctoral Research Project (JB24BYKY02) and the Key Research Project of Guangdong Province (2022B0202060005).

## Author contributions

H.-J.L. and H.W. designed and supervised the project. J.F. developed the software and performed the analysis. A.J. offered statistical advice. H.-J.L. and J.F. wrote the manuscript with revisions from all other authors.

---

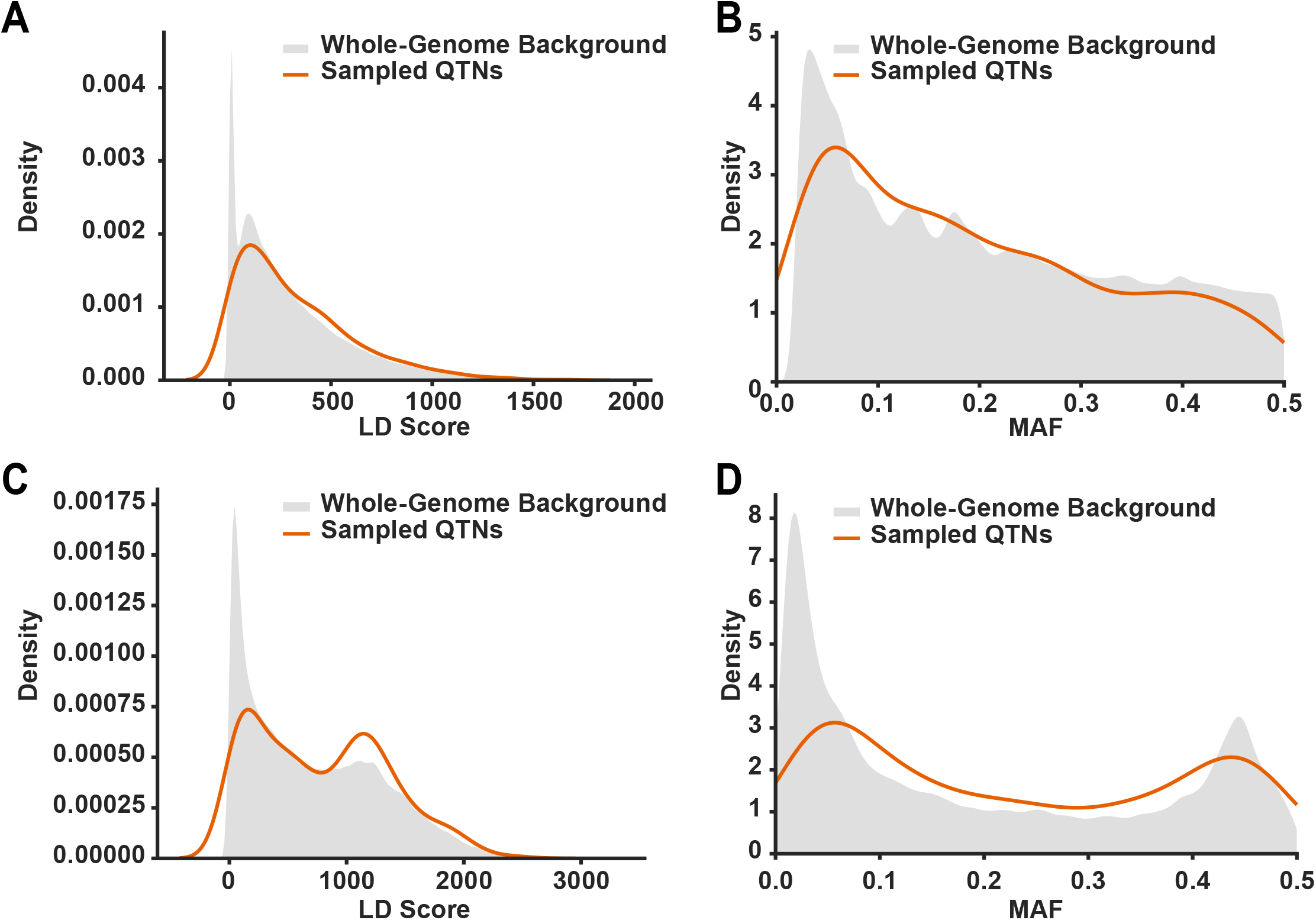

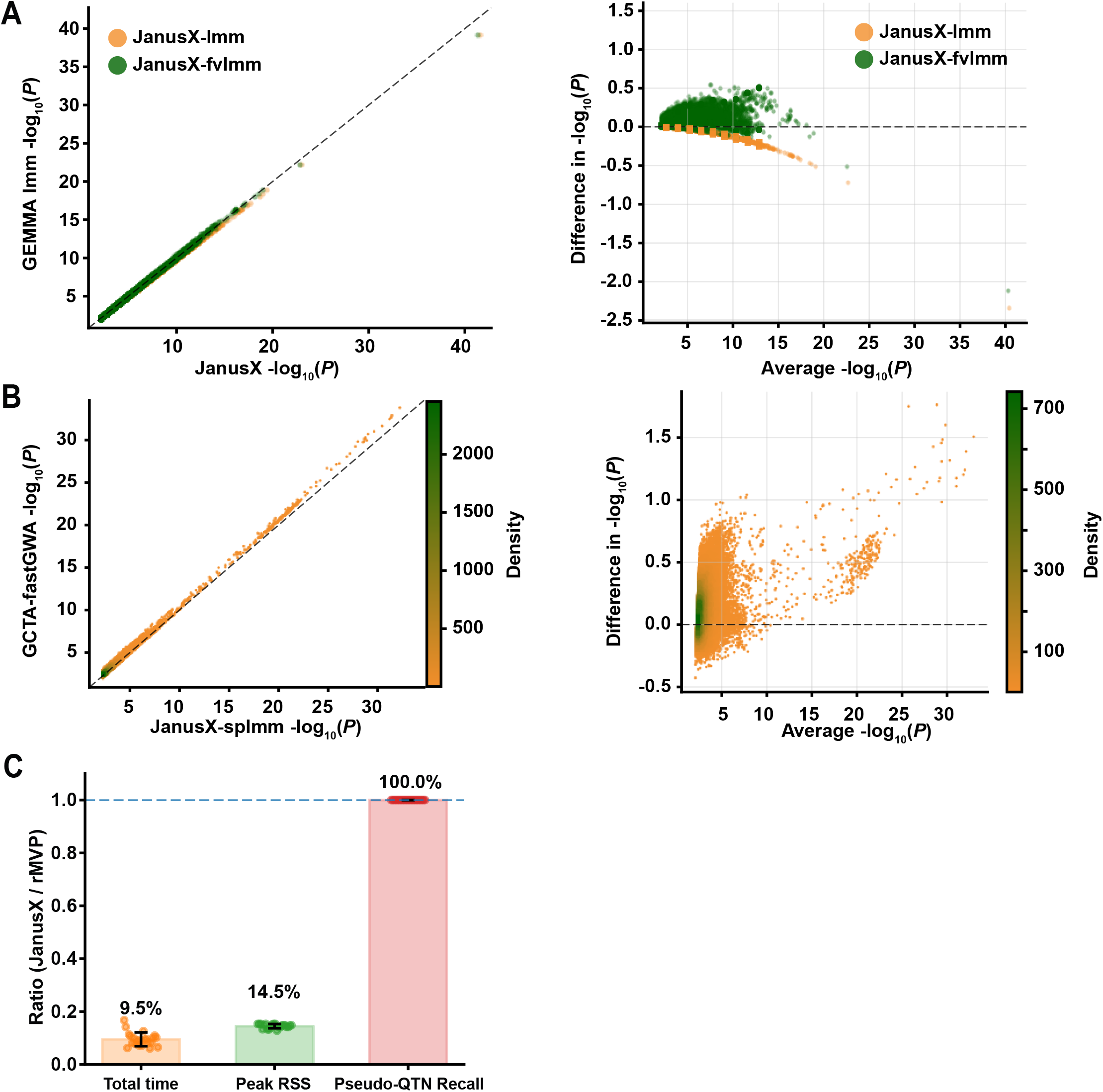

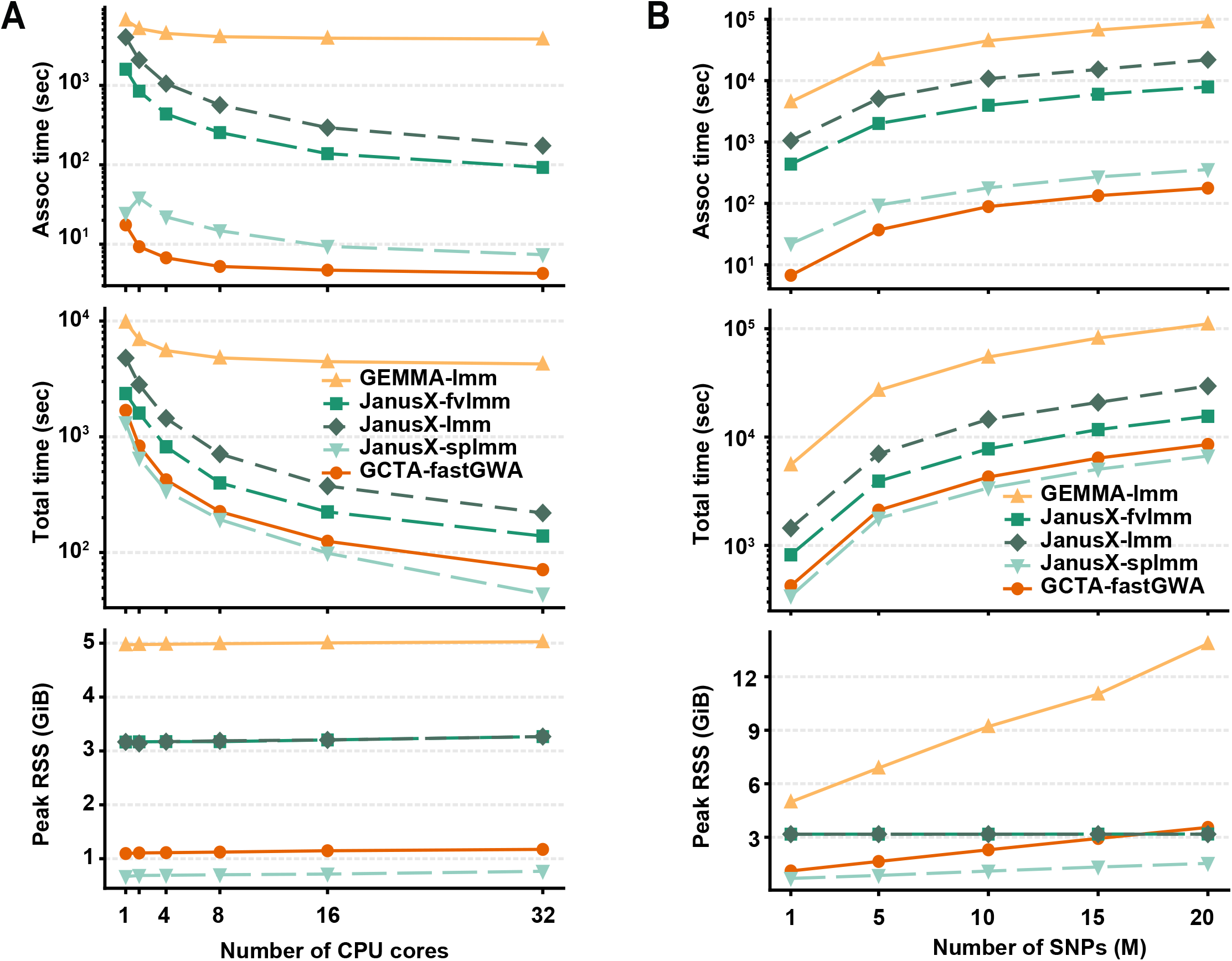

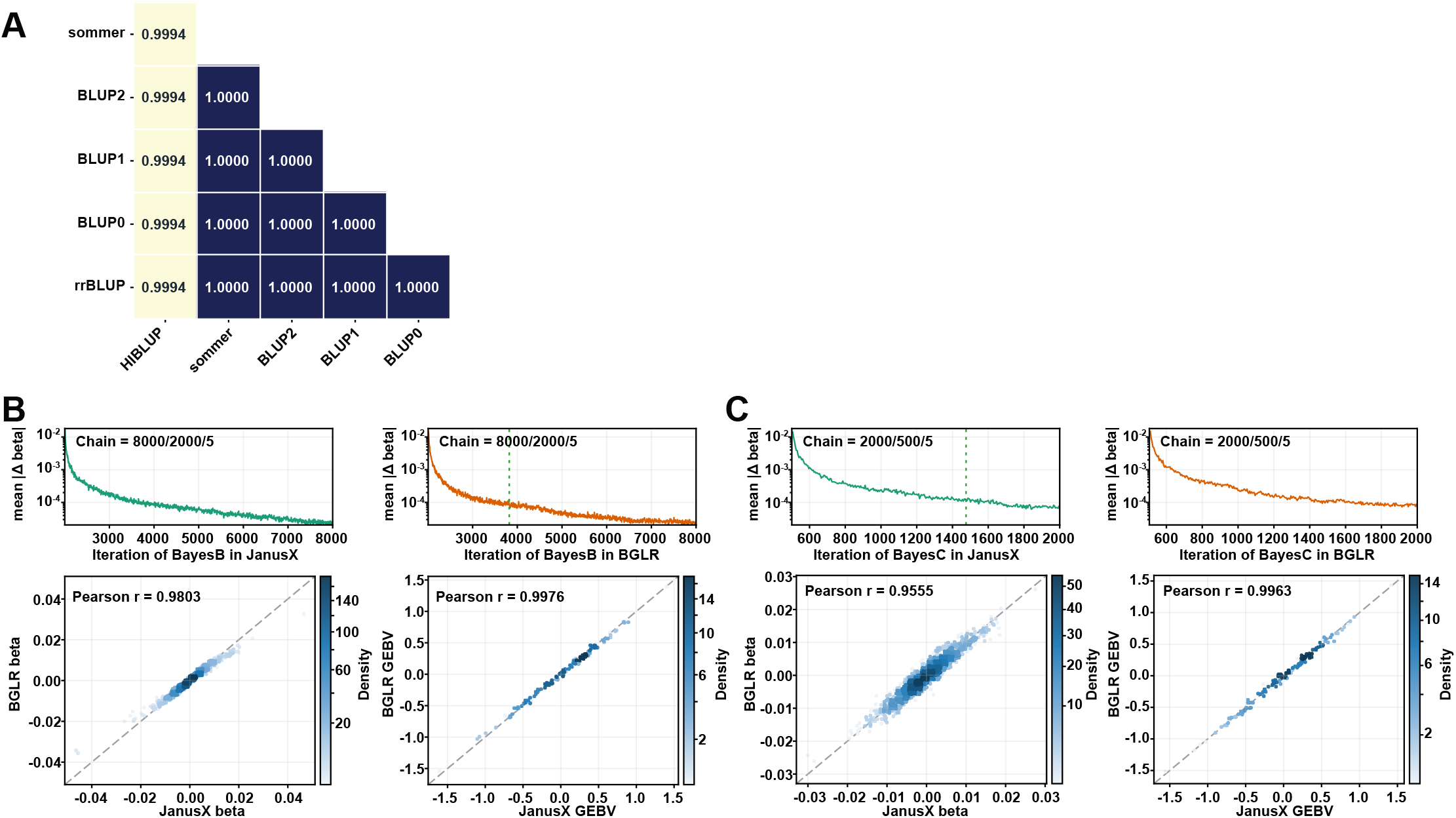

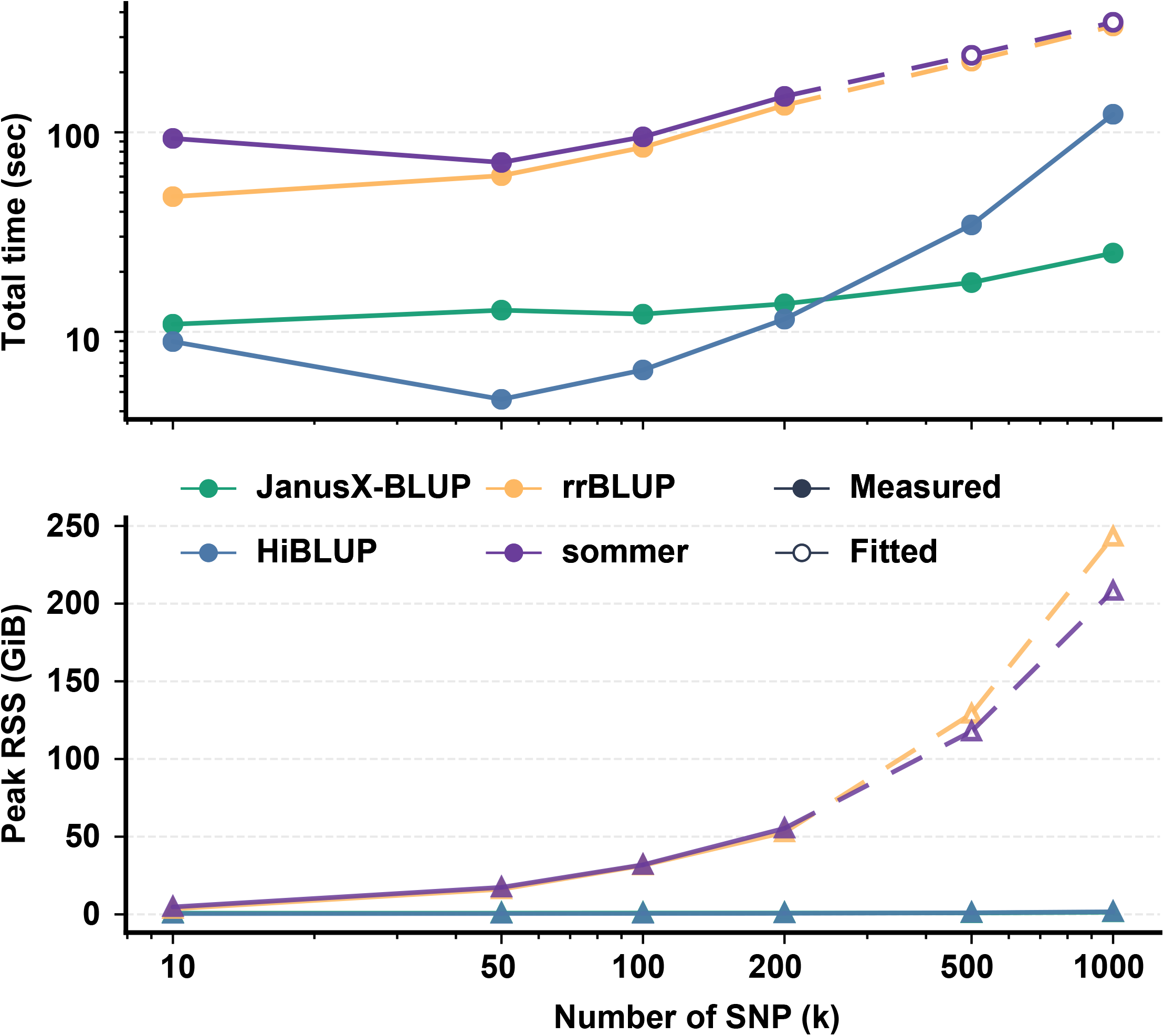

