## Supplemental Material for "JanusX: an integrated and high-performance platform for scalable genome-wide association studies and genomic selection"

### Supplemental information

Supplemental Note 1. **Visualization customization and reproducible figure generation in JanusX.**
Supplemental Note 2. **Algorithm in GWAS module.**
Supplemental Note 3. **BLUP-family algorithm in GS module.**
Supplemental Note 4. **Algorithm of Bayesian prediction models in GS module.**

Figure S1. **Distributions of Linkage Disequilibrium (LD) scores and Minor Allele Frequencies (MAF) for sampled causal variants.**
Figure S2. **Concordance and performance benchmarking of the JanusX GWAS module.**
Figure S3. **CPU scalability and SNP-size stress tests of JanusX GWAS backends.**
Figure S4. **Concordance of BLUP-family and Bayesian models in JanusX GS module.**
Figure S5. **SNP-size scalability of BLUP-family GS workflows.**

Table S1. **Simulation-based concordance of association statistics among JanusX-lmm (lmm), JanusX-fvlmm (fvlmm) and GEMMA-lmm (ref).**
Table S2. **Simulation-based concordance of association statistics between GCTA-fastGWA (ref) and JanusX-splmm (splmm).**
Table S3. **Three-way concordance of association statistics among GEMMA-lmm (ref), JanusX-lmm (lmm), and JanusX-fvlmm (fvlmm) in real maize CUBIC data.**
Table S4. **Two-way concordance of association statistics between GCTA-fastGWA (ref) and JanusX-splmm (splmm) in real maize data.**

### Supplemental Note 1. Visualization customization and reproducible figure generation in JanusX

JanusX provides dedicated post-analysis tools to ensure figure generation is explicit, customizable, and reproducible from the command line. Visualization functions are implemented for Genome-wide Association Studies (GWAS), genomic selection (GS), population-structure analysis, and Principal Components Analysis (PCA), and all the relevant graphical parameters can be specified directly by users rather than relying on fixed internal settings.

For GWAS post-processing, the `postgwas` module is used to generate Manhattan plots, quantile-quantile (QQ) plots, and Linkage Disequilibrium (LD) triangular block plots. Manhattan and QQ plots are generated from marker-level association results. Users can specify parameters including chromosome, physical-position, *P*-value columns, significance thresholds, genomic sub-ranges, y-axis limits, marker size, color palette, density-rendering mode, and output formats (PNG, PDF, SVG, and TIFF). For LD visualization, the `postgwas` module can generate triangular LD block plots based on genotype input, allowing users to control plot range, panel aspect ratio, and LD-specific color schemes.

For GS post-processing, the `postgs` module is used to generate model-evaluation plots and Manhattan-style marker-effect plots. Model-evaluation plots are produced from prediction outputs and observed phenotypes, with user-defined parameters such as figure format, color palette, panel aspect ratio, scatter-point size, and density-rendering mode. Marker-effect plots support user-specified input effect files, effect columns, signed effect visualization, marker-sizes, color palettes, and output formats. These options allow consistent analysis and visualization of prediction performance across BLUP-family, Bayesian, and machine-learning models.

For population-structure analysis, the `fastpop` module provides visualization of ancestry-proportion estimates as stacked bar plots. The number of ancestry components can be specified as a single value, range, or list of K values. Additional customization options include sample-label inputs for bar plots, as well as skipping plot generation for batch-mode output focus. Parameters such as optimizer selection, iteration limits, convergence tolerance, likelihood-check intervals, random seeds, thread counts, memory allocation, and optional cross-validation can also be explicitly modified to improve reproducibility in ancestry inference.

For PCA-related visualizations, the `pca` module accepts genotype or Genomic Relationship Matrix (GRM) input. Users can specify the output dimensions, variant-quality filters, grouping variables, color palettes, and settings for two-dimensional scatter-plots. Three-dimensional rotating animation visualizations in PC space (e.g. PC1-PC3) can also be generated to facilitate deeper insights into population structure where needed.

These customizable post-analysis modules make analytical and graphical settings explicit and reproducible across GWAS, GS, PCA, and ancestry analyses. By exposing parameters including input columns, plotting ranges, point sizes, transparency, density-rendering modes, color palettes, reference-thresholds, figure formats, and output resolutions, JanusX allows users to generate high-quality, publication-ready figure without compromising command-line reproducibility.

### Supplemental Note 2. Algorithm in GWAS module

In the linear model, we define a phenotype vector $\mathbf{y} \in \mathbb{R}^n$, covariate matrix $\mathbf{X} \in \mathbb{R}^{n\times p}$, and marker vector $\mathbf{g}_j \in \mathbb{R}^n$. Then, JanusX-lm fits the fixed-effect marker model:

$$\mathbf{y} = \mathbf{X\beta} + \mathbf{g}_j\boldsymbol{\alpha}_j + \boldsymbol{\varepsilon}, \qquad \boldsymbol{\varepsilon} \sim \mathcal{N}\left(\mathbf{0}, \sigma_\varepsilon^2\mathbf{I}\right),$$

where $\boldsymbol{\beta}$ represents covariate effects, $\boldsymbol{\alpha}_j$ is the marker effect, and ε is the residual term. To avoid repeated covariate fitting across markers, JanusX applies the Frisch-Waugh-Lovell projection:

$$\mathbf{P}_X = \mathbf{I} - \mathbf{X}\left(\mathbf{X}^T\mathbf{X}\right)^{-1}\mathbf{X}^T, \qquad \tilde{\mathbf{y}} = \mathbf{P}_X\mathbf{y}, \qquad \tilde{\mathbf{g}}_j = \mathbf{P}_X\mathbf{g}_j.$$

This approach reduces marker-level analysis to univariate regression of transformed data ($\tilde{\mathbf{y}}$ on $\tilde{\mathbf{g}}_j$), avoiding repetitive covariate remodeling.

In the linear mixed model, JanusX-lmm incorporates a random additive genetic effect ($\mathbf{u}$):

$$\mathbf{y} = \mathbf{X\beta} + \mathbf{g}_j\boldsymbol{\alpha}_j + \mathbf{u} + \boldsymbol{\varepsilon}, \qquad \mathbf{u} \sim \mathcal{N}\left(\mathbf{0}, \sigma_g^2\mathbf{K}\right), \qquad \boldsymbol{\varepsilon} \sim \mathcal{N}\left(\mathbf{0}, \sigma_\varepsilon^2\mathbf{I}\right),$$

where $\mathbf{K}$ is the genomic relationship matrix (GRM) and

$$\mathbf{V} = \sigma_g^2\mathbf{K} + \sigma_\varepsilon^2\mathbf{I}.$$

Using EVD,

$$\mathbf{K} = \mathbf{U\Lambda U}^T,$$

followed by transformation of $\mathbf{y}$, $\mathbf{X}$, and $\mathbf{g}_j$ into the eigenvector space, JanusX-lmm performs efficient mixed-model fitting in diagonalized covariance matrices to reduce computation time for genome-wide scans, whereas JanusX-fvlmm estimates variance components once under the null model to ensure much more efficient computation.

For large-cohort analysis, JanusX-splmm reduces computational demand by creating a sparse-GRM:

$$K_{ij}^{(s)} = K_{ij} \cdot \mathbf{1}\left(\left|K_{ij}\right| \geq \tau\right), \qquad i \neq j.$$

Marker testing based on null-model residuals is calibrated using a genome-wide GRAMMAR-Gamma calibration factor ($\gamma$) estimated from 1,000 randomly sampled SNPs, providing rapid approximate association scans when dense-GRM LMM inference is computationally infeasible.

JanusX also implements FarmCPU, alternating between fixed-effect marker testing and random-effect pseudo-QTN optimization. Through parallelized model fitting and packed genotype processing, JanusX improves FarmCPU's computational speed while maintaining its statistical framework.

### Supplemental Note 3. BLUP-family algorithm in GS module

For GBLUP models, JanusX uses

$$\boldsymbol{y} = \boldsymbol{Xb} + \boldsymbol{g} + \boldsymbol{\varepsilon}, \qquad \boldsymbol{g} \sim \mathcal{N}\left(\mathbf{0}, \sigma_g^2\boldsymbol{K}\right), \qquad \boldsymbol{e} \sim \mathcal{N}\left(\mathbf{0}, \sigma_\varepsilon^2\boldsymbol{I}\right),$$

where $K$ is the Genomic Relationship Matrix (GRM) derived from the centered marker matrix $W$ and scaled marker-count constant $c$:

$$\boldsymbol{K} = \frac{\boldsymbol{W}\boldsymbol{W}^{\boldsymbol{T}}}{\boldsymbol{c}}.$$

In the default JanusX BLUP workflow, markers are centered using the same sample set used for the corresponding GRM construction, and the global scaling constant is matched across GBLUP, exact rrBLUP, and rrBLUP-PCG to preserve solver-space concordance. Let

$$\boldsymbol{V} = \boldsymbol{\sigma}_{\boldsymbol{g}}^{2}\boldsymbol{K} + \boldsymbol{\sigma}_{\boldsymbol{\varepsilon}}^{2}\boldsymbol{I} = \boldsymbol{\sigma}_{\boldsymbol{g}}^{2}(\boldsymbol{K} + \boldsymbol{\lambda}\boldsymbol{I}), \qquad \boldsymbol{\lambda} = \frac{\boldsymbol{\sigma}_{\boldsymbol{\varepsilon}}^{2}}{\boldsymbol{\sigma}_{\boldsymbol{g}}^{2}}.$$

After estimating variance components, the GBLUP solution can be expressed as

$$\widehat{\boldsymbol{g}} = \boldsymbol{K}(\boldsymbol{K} + \boldsymbol{\lambda}\boldsymbol{I})^{-1}(\boldsymbol{y} - \boldsymbol{X}\widehat{\boldsymbol{b}}).$$

For held-out samples, prediction uses the cross-kernel between test and training samples,

$$\widehat{\boldsymbol{g}}_{*} = \boldsymbol{K}_{*\boldsymbol{t}}(\boldsymbol{K}_{\boldsymbol{tt}} + \boldsymbol{\lambda}\boldsymbol{I})^{-1}(\boldsymbol{y}_{\boldsymbol{t}} - \boldsymbol{X}_{\boldsymbol{t}}\widehat{\boldsymbol{b}}),$$

where $K_{tt}$ is the training-sample GRM and $K_{*t}$ is the test-by-training cross-kernel. To minimize memory usage, block-wise GRM streaming is used instead of full dense matrix storage. In cross-validation settings, when memory allows, JanusX can construct a shared full-sample GRM once and reuse train/train and test/train submatrices across folds. For moderate sample sizes, variance components are estimated after spectral decomposition of the training GRM, providing a computationally efficient workflow.

When the number of individuals becomes large relative to the number of effective markers, JanusX can switch from sample-space GBLUP to marker-space rrBLUP to avoid the cubic cost of decomposing an $n \times n$ matrix. In marker space, the ridge-regression equation:

$$(\boldsymbol{W}^{\boldsymbol{T}}\boldsymbol{M}_{\boldsymbol{X}}\boldsymbol{W} + \boldsymbol{\lambda}\boldsymbol{I}_{\boldsymbol{m}})\widehat{\boldsymbol{\beta}} = \boldsymbol{W}^{\boldsymbol{T}}\boldsymbol{M}_{\boldsymbol{X}}\boldsymbol{y}, \qquad \boldsymbol{\lambda} = \frac{\boldsymbol{\sigma}_{\boldsymbol{\varepsilon}}^{2}}{\boldsymbol{\sigma}_{\boldsymbol{\beta}}^{2}}$$

is solved, where $M_X = I - X(X^TX)^{-1}X^T$ removes confounding effects, and the rrBLUP model estimates additive marker effects directly:

$$\boldsymbol{y} = \boldsymbol{X}\boldsymbol{b} + \boldsymbol{W}\boldsymbol{\beta} + \boldsymbol{\varepsilon}, \qquad \boldsymbol{\beta} \sim \boldsymbol{N}(\boldsymbol{0}, \boldsymbol{I}_{\boldsymbol{m}}\boldsymbol{\sigma}_{\boldsymbol{\beta}}^{2}), \qquad \boldsymbol{e} \sim \boldsymbol{N}(\boldsymbol{0}, \boldsymbol{I}\boldsymbol{\sigma}_{\boldsymbol{\varepsilon}}^{2}).$$

For the intercept-only case, this reduces to centering both phenotypes and markers before solving the ridge system. In exact rrBLUP, JanusX builds the marker-space Gram matrix

$$\boldsymbol{A} = \boldsymbol{W}^{\boldsymbol{T}}\boldsymbol{M}_{\boldsymbol{X}}\boldsymbol{W}$$

and performs spectral decomposition

$$\boldsymbol{A} = \boldsymbol{U}\mathbf{diag}(\boldsymbol{s})\boldsymbol{U}^{\boldsymbol{T}}.$$

Variance-component estimation is then performed in the non-zero spectral space, and the final marker effects are recovered after optimizing $\lambda$. This route is efficient when the number of effective markers is smaller than the number of individuals, because the dominant decomposition is performed in marker space rather than sample space. For large marker sets, explicitly constructing $W^TM_XW$ is memory- and time-intensive. JanusX therefore implements a matrix-free rrBLUP-PCG backend that solves

$$(\boldsymbol{W}^{\boldsymbol{T}}\boldsymbol{M}_{\boldsymbol{X}}\boldsymbol{W} + \boldsymbol{\lambda}\boldsymbol{I}_{\boldsymbol{m}})\widehat{\boldsymbol{\beta}} = \boldsymbol{W}^{\boldsymbol{T}}\boldsymbol{M}_{\boldsymbol{X}}\boldsymbol{y}$$

using the preconditioned conjugate gradient (PCG) algorithm. Each PCG iteration only requires the matrix-vector product

$$\boldsymbol{v} \mapsto \boldsymbol{W}^{\boldsymbol{T}} \boldsymbol{M}_{\boldsymbol{X}}(\boldsymbol{W}\boldsymbol{v}) + \boldsymbol{\lambda}\boldsymbol{v},$$

which can be computed by streaming genotype blocks from packed input. This avoids storing the full $m \times m$ marker Gram matrix and keeps memory usage bounded by block size, phenotype/covariate arrays, and solver vectors.

### Supplemental Note 4. Algorithm of Bayesian prediction models in GS module

JanusX currently implements BayesA, BayesB, and BayesC using Gibbs-sampling-based posterior inference:

$$\boldsymbol{y} = \boldsymbol{X}\boldsymbol{\alpha} + \boldsymbol{Z}\boldsymbol{\beta} + \boldsymbol{\varepsilon}, \qquad \boldsymbol{e} \sim \boldsymbol{N}\left(\mathbf{0}, \boldsymbol{\sigma}_{\boldsymbol{\varepsilon}}^{2}\boldsymbol{I}\right),$$

where $Z$ is the standardized marker matrix and $\beta$ is the vector of marker effects.
In BayesA, each marker has an independent variance prior:

$$\boldsymbol{\beta}_{\boldsymbol{j}} \mid \boldsymbol{\sigma}_{\boldsymbol{\beta j}}^{2} \sim \boldsymbol{N}\left(\mathbf{0}, \boldsymbol{\sigma}_{\boldsymbol{\beta j}}^{2}\right), \qquad \boldsymbol{\sigma}_{\boldsymbol{\beta j}}^{2} \sim \textbf{scaled-inv-}\boldsymbol{\chi}^{\mathbf{2}},$$

allowing marker-specific shrinkage. BayesB/C introduce a marker inclusion indicator which is determined by Bernoulli trials:

$$\boldsymbol{\delta}_{\boldsymbol{j}} \sim \textbf{Bernoulli}(\boldsymbol{\pi}), \qquad \boldsymbol{\beta}_{\boldsymbol{j}} = \mathbf{0} \textbf{ if } \boldsymbol{\delta}_{\boldsymbol{j}} = \mathbf{0},$$

where active markers follow a marker-specific variance prior. BayesC also uses a spike-and-slab prior, and assigns a shared variance for active markers,

$$\boldsymbol{\beta}_{\boldsymbol{j}} \mid \boldsymbol{\delta}_{\boldsymbol{j}} = \mathbf{1} \sim \boldsymbol{N}\left(\mathbf{0}, \boldsymbol{\sigma}_{\boldsymbol{\beta}}^{2}\right),$$

and the inclusion probability $\pi$ is updated from its posterior distribution. For posterior prediction, retained MCMC samples after burn-in and thinning are averaged to obtain posterior mean effects. Test-sample predictions are computed as

$$\widehat{\boldsymbol{y}}_{*} = \boldsymbol{X}_{*}\bar{\boldsymbol{\alpha}} + \boldsymbol{Z}_{*}\bar{\boldsymbol{\beta}},$$

where $\bar{\alpha}$ and $\bar{\beta}$ are posterior means over retained samples, ensuring predictions align with Bayesian posterior distributions.

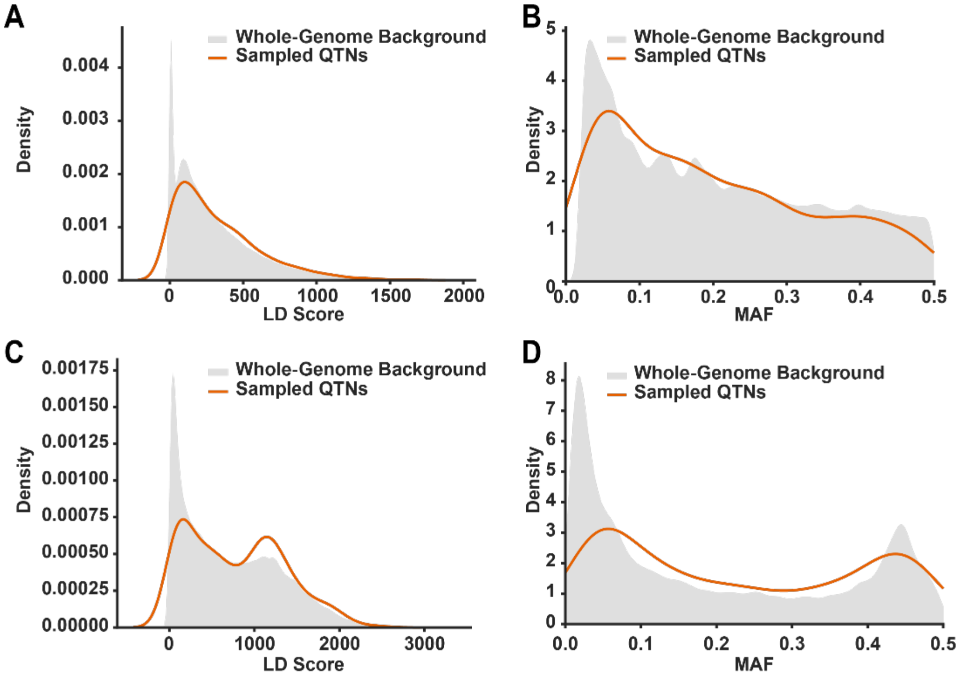


**Figure S1. Distributions of Linkage Disequilibrium (LD) scores and Minor Allele Frequencies (MAF) for sampled causal variants. (A, B)** Density plots comparing whole-genome LD scores and MAF distributions to those of randomly sampled variants in the maize CUBIC population, which included 1,493 individuals and 8,666,018 SNPs. **(C, D)** Similar density comparisons for LD scores and MAF in the rice population, which contained 3,750 individuals and 4,855,965 SNPs. The grey shaded areas and red curves denote the whole-genome background and distributions of randomly sampled variants, respectively.

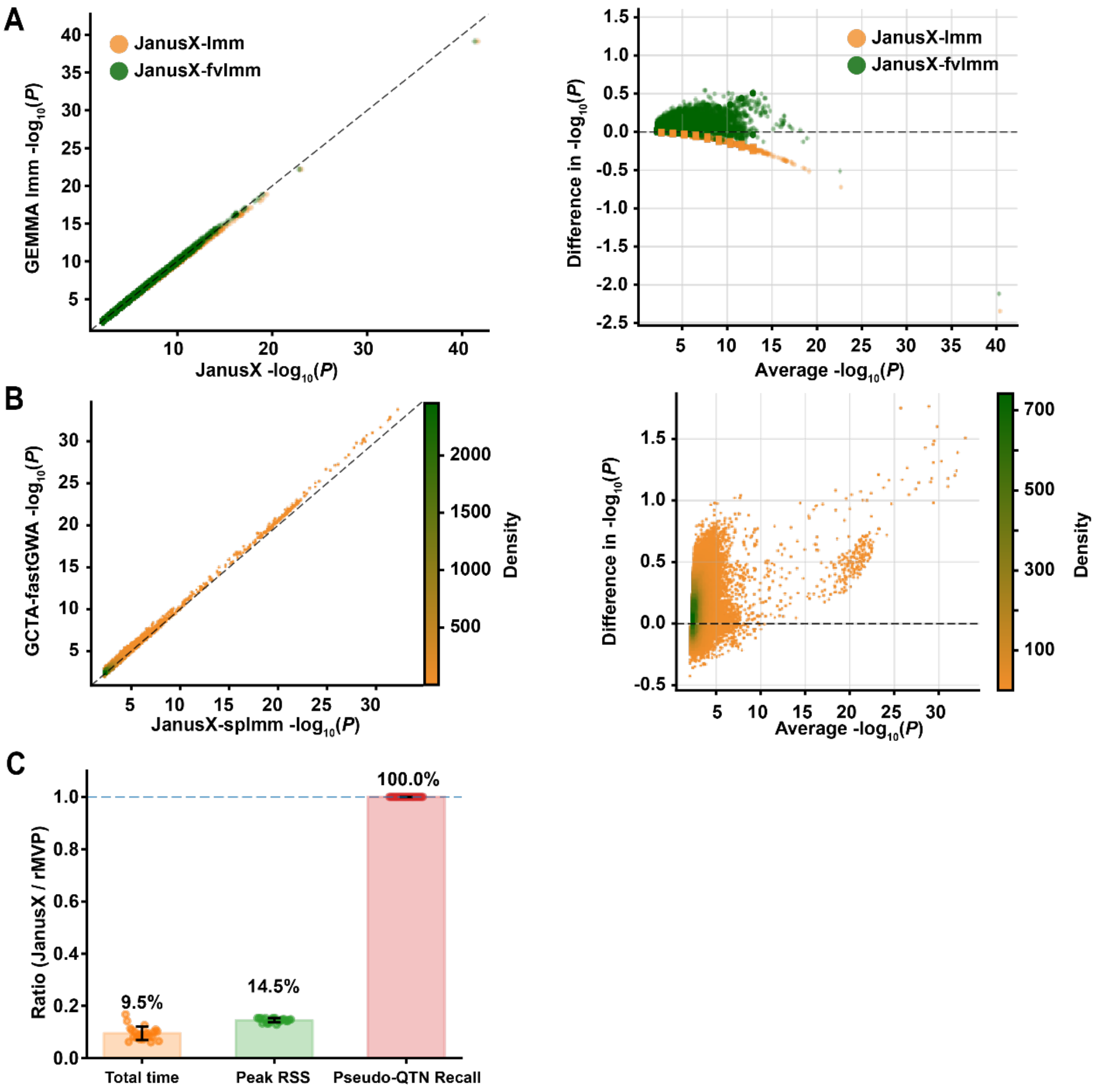


**Figure S2. Concordance and performance benchmarking of the JanusX GWAS module. (A, B)** Concordance of JanusX GWAS implementations with reference tools in representative simulation benchmarks generated from empirical maize CUBIC genotype matrices. Scatter plots show agreement in association significance among the top 1% most significant SNPs, with the black dashed diagonal line indicating perfect concordance (x = y). Bland-Altman plots show the corresponding agreement pattern for the same SNP set, with differences calculated as reference-tool values minus JanusX values. (A) Concordance of JanusX-lmm and JanusX-fvlmm with GEMMA-lmm. Yellow and green points represent JanusX-lmm and JanusX-fvlmm, respectively. (B) Concordance of JanusX-splmm with GCTA-fastGWA. Points are color-coded by density, with yellow for low density and green for high density. Both panels use benchmarks simulated from empirical maize CUBIC genotypes, with 10 causal QTNs and phenotypic variance partitioning of $V_{QTN}$:$V_{Bg}$:$V_E$ = 0.3:0.3:0.4. **(C)** Performance comparison of pseudo-QTN concordance between JanusX-FarmCPU and rMVP-FarmCPU across 23 agronomic traits in the maize CUBIC panel. Bars show the JanusX/rMVP runtime and peak memory usage ratios, together with the pseudo-QTN recall. RSS denotes resident set size.

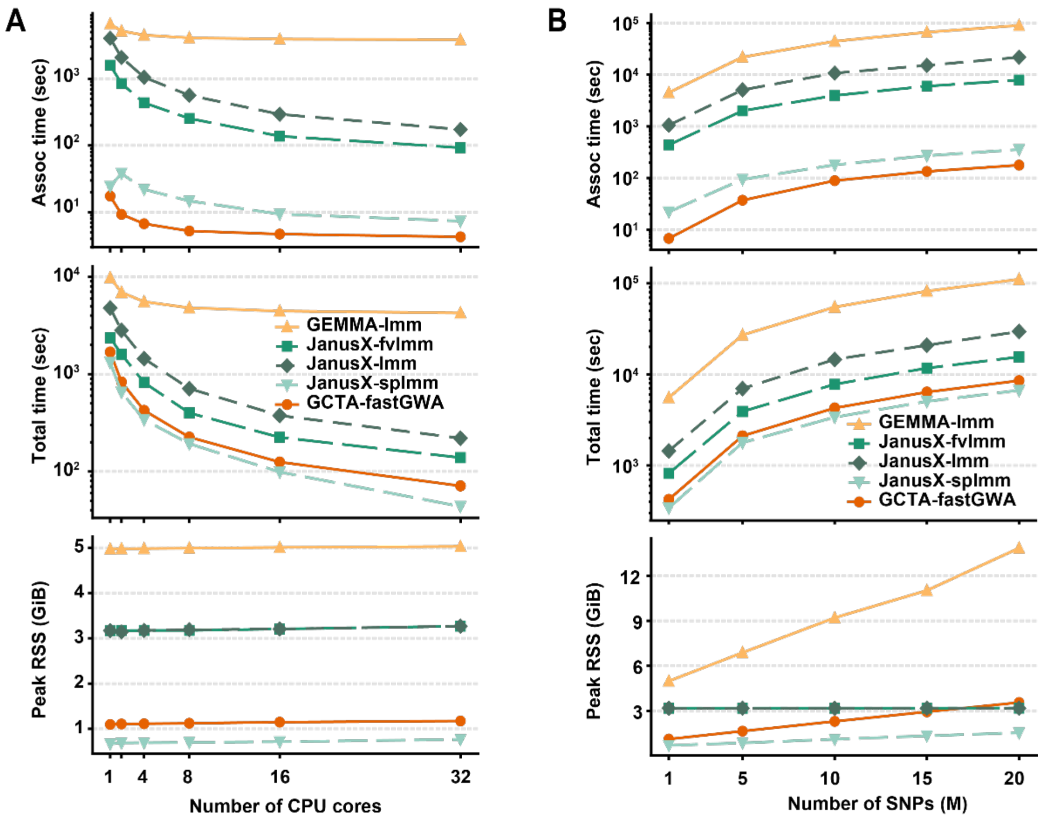


**Figure S3. CPU scalability and SNP-size stress tests of JanusX GWAS backends. (A)** CPU scalability for an analysis with 10,000 individuals and 1,000,000 SNPs across 1-32 CPU cores. **(B)** SNP-size scalability benchmark using 10,000 individuals with variable marker counts. For both tests, runtime at the association-stage (Assoc. Time), end-to-end runtime (including GRM construction), and peak resident set size (RSS) were measured. GiB, gibibyte.

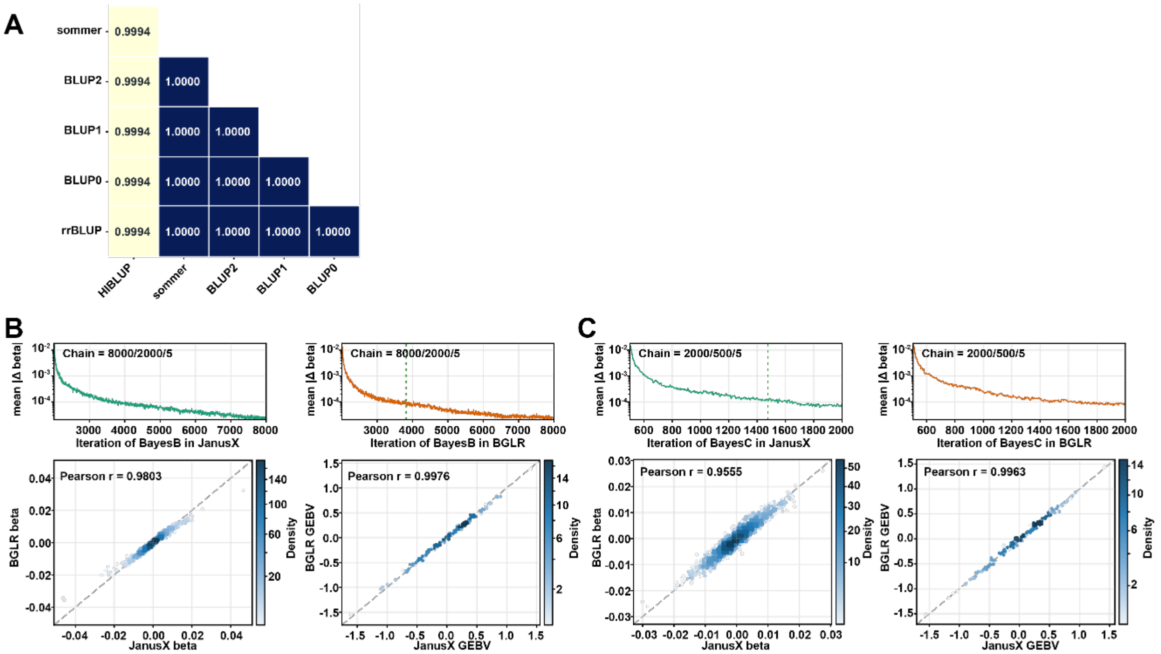


**Figure S4. Concordance of BLUP-family and Bayesian models in JanusX GS module.** This benchmark is evaluated using the wheat benchmark dataset distributed in the BGLR package. **(A)** A heatmap showing GEBV concordance between JanusX BLUP-family models and reference tools (sommer, rrBLUP, HIBLUP). BLUP0-2 are BLUP-family models in JanusX: BLUP0 for GBLUP, BLUP1 for rrBLUP, and BLUP2 for rrBLUP with Preconditioned Conjugate Gradient (PCG) solver. All methods employed identical SNP centering strategies. **(B, C)** Comparison of Bayesian models between JanusX and BGLR using convergence diagnostics and Pearson correlation analyses. Trace plots show the convergence of marker-effect (beta) for BayesB and BayesC. Scatter plots compare posterior SNP marker effects and GEBVs, with Pearson correlation coefficients (r) quantifying concordance. MCMC parameters include total iterations, burn-in, and thinning intervals. Point density is color-coded.

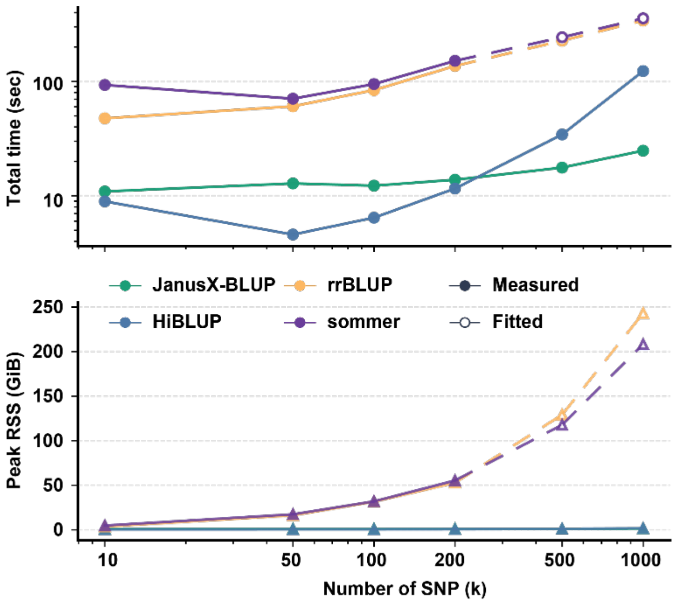


**Figure S5. SNP-size scalability of BLUP-family GS workflows.** Comparison of total runtime and peak RSS across increasing cohort sizes, tested with 5k individuals, between JanusX (GBLUP from GS module) and reference tools (HIBLUP, sommer, rrBLUP). All methods applied consistent marker filters and 5-fold cross-validation splits. Uncompleted runs result from memory usage exceeding 50 GiB. Measured values and model-fitted estimates are shown. RSS denotes resident set size; GiB denotes gibibyte.

**Table S1. Simulation-based concordance of association statistics among JanusX-lmm (lmm), JanusX-fvlmm (fvlmm) and GEMMA-lmm (ref).** Concordance was evaluated using the top 1% most significant SNPs from simulated phenotypes for maize CUBIC and RiceAtlas genotypes across diverse genetic-architecture scenarios. $\lambda_{GC}$ values represent the mean across 10 replicates. Pearson correlations (Pearson r) were calculated using estimated marker effects (beta), and Spearman correlations (Spearman ρ) were calculated using association significance. QTN denotes Quantitative Trait Nucleotide. $V_{QTN}$:$V_{Bg}$:$V_E$ denotes the simulated phenotypic variance partitioned into causal-QTN variance, polygenic background variance, and residual environmental variance, respectively.

| Dataset | QTNs | Variance ($V_{QTN}$:$V_{Bg}$:$V_E$) | lmm Pearson r (beta) | lmm Spearman ρ ($-\log_{10}P$) | fvlmm Pearson r (beta) | fvlmm Spearman ρ ($-\log_{10}P$) | $\lambda_{GC}$ (ref / lmm / fvlmm) |
|---|---|---|---|---|---|---|---|
| RiceAtlas | 10 | 0.5:0.0:0.5 | 1.0000 ± 0.0000 | 1.0000 ± 0.0000 | 0.9999 ± 0.0001 | 0.9962 ± 0.0052 | 0.763 / 0.763 / 0.762 |
| | 10 | 0.3:0.3:0.4 | 1.0000 ± 0.0000 | 1.0000 ± 0.0000 | 0.9999 ± 0.0001 | 0.9982 ± 0.0025 | 0.868 / 0.868 / 0.867 |
| | 10 | 0.1:0.6:0.3 | 1.0000 ± 0.0000 | 1.0000 ± 0.0000 | 1.0000 ± 0.0000 | 0.9983 ± 0.0018 | 0.951 / 0.951 / 0.950 |
| | 100 | 0.5:0.0:0.5 | 1.0000 ± 0.0000 | 1.0000 ± 0.0000 | 1.0000 ± 0.0000 | 0.9976 ± 0.0026 | 0.896 / 0.896 / 0.895 |
| | 100 | 0.3:0.3:0.4 | 1.0000 ± 0.0000 | 1.0000 ± 0.0000 | 1.0000 ± 0.0000 | 0.9983 ± 0.0021 | 0.943 / 0.943 / 0.942 |
| | 100 | 0.1:0.6:0.3 | 1.0000 ± 0.0000 | 1.0000 ± 0.0000 | 1.0000 ± 0.0000 | 0.9990 ± 0.0006 | 0.985 / 0.985 / 0.984 |
| CUBIC | 10 | 0.5:0.0:0.5 | 0.9995 ± 0.0005 | 1.0000 ± 0.0000 | 0.9991 ± 0.0006 | 0.9844 ± 0.0058 | 0.867 / 0.868 / 0.865 |
| | 10 | 0.3:0.3:0.4 | 0.9997 ± 0.0003 | 1.0000 ± 0.0000 | 0.9996 ± 0.0003 | 0.9929 ± 0.0035 | 0.962 / 0.962 / 0.960 |
| | 10 | 0.1:0.6:0.3 | 0.9998 ± 0.0001 | 1.0000 ± 0.0000 | 0.9998 ± 0.0001 | 0.9982 ± 0.0007 | 0.998 / 0.999 / 0.996 |
| | 100 | 0.5:0.0:0.5 | 0.9998 ± 0.0001 | 1.0000 ± 0.0000 | 0.9998 ± 0.0001 | 0.9948 ± 0.0026 | 0.984 / 0.984 / 0.981 |
| | 100 | 0.3:0.3:0.4 | 0.9998 ± 0.0001 | 1.0000 ± 0.0000 | 0.9998 ± 0.0001 | 0.9975 ± 0.0009 | 1.003 / 1.004 / 1.001 |
| | 100 | 0.1:0.6:0.3 | 0.9998 ± 0.0001 | 1.0000 ± 0.0000 | 0.9998 ± 0.0001 | 0.9983 ± 0.0010 | 1.005 / 1.006 / 1.003 |

**Table S2. Simulation-based concordance of association statistics between GCTA-fastGWA (ref) and JanusX-splmm (splmm)**. Performance was evaluated using the top 1% most significant SNPs. Benchmarks used simulated phenotypes from maize CUBIC and RiceAtlas genotype data.

| Dataset | QTNs | Variance ($V_{QTN}$:$V_{Bg}$:$V_E$) | Pearson r (beta) | Spearman ρ (-log$_{10}$*P*) | $\lambda_{GC}$ (ref / splmm) |
|---|---|---|---|---|---|
| RiceAtlas | 10 | 0.5:0.0:0.5 | 1.0000 ± 0.0000 | 0.9998 ± 0.0001 | 0.450 / 0.422 |
| | 10 | 0.3:0.3:0.4 | 1.0000 ± 0.0000 | 0.9988 ± 0.0022 | 0.489 / 0.456 |
| | 10 | 0.1:0.6:0.3 | 1.0000 ± 0.0001 | 0.9967 ± 0.0043 | 0.465 / 0.430 |
| | 100 | 0.5:0.0:0.5 | 1.0000 ± 0.0000 | 0.9996 ± 0.0001 | 0.521 / 0.486 |
| | 100 | 0.3:0.3:0.4 | 1.0000 ± 0.0000 | 0.9992 ± 0.0008 | 0.581 / 0.536 |
| | 100 | 0.1:0.6:0.3 | 1.0000 ± 0.0000 | 0.9972 ± 0.0032 | 0.494 / 0.456 |
| CUBIC | 10 | 0.5:0.0:0.5 | 1.0000 ± 0.0000 | 1.0000 ± 0.0000 | 0.746 / 0.747 |
| | 10 | 0.3:0.3:0.4 | 1.0000 ± 0.0000 | 1.0000 ± 0.0000 | 0.855 / 0.842 |
| | 10 | 0.1:0.6:0.3 | 0.9998 ± 0.0007 | 0.9804 ± 0.0589 | 0.914 / 0.892 |
| | 100 | 0.5:0.0:0.5 | 1.0000 ± 0.0000 | 1.0000 ± 0.0000 | 0.964 / 0.945 |
| | 100 | 0.3:0.3:0.4 | 1.0000 ± 0.0000 | 1.0000 ± 0.0000 | 0.927 / 0.906 |
| | 100 | 0.1:0.6:0.3 | 1.0000 ± 0.0000 | 1.0000 ± 0.0000 | 0.933 / 0.915 |

**Table S3. Three-way concordance of association statistics among GEMMA-lmm (ref), JanusX-lmm (lmm), and JanusX-fvlmm (fvlmm) in real maize CUBIC data.** Concordance was evaluated using the top 1% most significant SNPs across 5 representative agronomic traits. Pearson r was calculated using estimated marker effects (beta), and Spearman ρ was calculated using association significance. Traits include DTA (days to anthesis), EH (ear height), PH (plant height), EW (ear weight), and KNPE (kernel number per ear).

| Trait | lmm Pearson r (beta) | lmm Spearman ρ ($-\log_{10}P$) | fvlmm Pearson r (beta) | fvlmm Spearman ρ ($-\log_{10}P$) | $\lambda_{GC}$ (ref / lmm / fvlmm) |
|---|---|---|---|---|---|
| **DTA** | 1.0000 | 1.0000 | 1.0000 | 1.0000 | 0.8974 / 0.8959 / 0.8936 |
| **EH** | 1.0000 | 1.0000 | 1.0000 | 1.0000 | 0.9248 / 0.9241 / 0.9213 |
| **PH** | 1.0000 | 1.0000 | 1.0000 | 1.0000 | 0.9380 / 0.9358 / 0.9330 |
| **EW** | 1.0000 | 1.0000 | 1.0000 | 1.0000 | 0.9554 / 0.9537 / 0.9500 |
| **KNPE** | 1.0000 | 1.0000 | 1.0000 | 1.0000 | 0.9576 / 0.9558 / 0.9520 |

**Table S4. Two-way concordance of association statistics between GCTA-fastGWA (ref) and JanusX-splmm (splmm) in real maize data.** Top 1% SNP concordance assessed using Pearson r (marker effects beta) and Spearman ρ (association significance) across DTA, EH, PH, EW, and KNPE.

| Trait | Pearson r (beta) | Spearman ρ ($-\log_{10}P$) | $\lambda_{GC}$ (ref / splmm) |
|---|---|---|---|
| **DTA** | 0.9993 | 0.9985 | 0.8273 / 0.8254 |
| **EH** | 0.9996 | 0.9993 | 0.9651 / 0.9640 |
| **PH** | 0.9996 | 0.9991 | 0.8678 / 0.8646 |
| **EW** | 0.9998 | 0.9995 | 0.9070 / 0.8832 |
| **KNPE** | 0.9998 | 0.9997 | 0.9993 / 0.9851 |
